# Heart Rate Variability with Photoplethysmography in 8 Million Individuals: Results and Scaling Relations with Age, Gender, and Time of Day

**DOI:** 10.1101/772285

**Authors:** Aravind Natarajan, Alexandros Pantelopoulos, Hulya Emir-Farinas, Pradeep Natarajan

## Abstract

Heart rate variability, or the variation in the time interval between consecutive beats, is a non-invasive dynamic metric of the autonomic nervous system and an independent risk factor for cardiovascular death. Prior limitations of use include requirements for continuous electrocardiography and lack of reference standards. Consumer wrist-worn tracking devices using photoplethysmography now provide the unique potential of continuously measuring surrogates of sympathetic and parasympathetic activity through the analysis of interbeat intervals. Here we leverage wrist-worn trackers to present the largest, to our knowledge, analysis of heart rate variability in humans across the time, frequency, and graphical domains. We derive diurnal parasympathetic and sympathetic measures and provide scaling parameters by age, sex, and time of day. Poincare plots graphically summarize heart rate variability metrics and may detect common arrhythmias. Lastly, we observe a strong dose-dependent correlation between daily steps and optimal heart rate variability metrics. Our results provide the ability to interpret continuous heart rate variability for tens of millions of wrist-worn trackers already in use.

## Introduction

Heart rate variability (HRV) refers to the variation in time between successive heart beats, and represents a non-invasive index of the autonomic nervous system. Since the autonomic nervous system regulates heart rate during sinus rhythm, HRV summarizes complex non-linear cardiovascular accommodative responses, dictated by the parasympathetic and sympathetic nervous systems, to dynamic physiologic variations.

While HRV is significantly influenced by sex and aging (*1*), reduced compensatory response (i.e., low HRV) is independently predictive of first fatal and non-fatal cardiovascular disease events in the general population (*2–6*). Robust data also links low HRV with adverse outcomes and mortality after sustaining a cardiovascular event, such as myocardial infarction (*7, 8*). Beta-blockers and exercise therapy reduce risks of cardiovascular events among individuals with coronary artery disease and congestive heart failure, and enhancement of HRV is believed to be mechanisms for improved prognosis (*9–11*). Thus, a less adaptive autonomic nervous system is predictive of first and recurrent cardiovascular events, and restoration of homeostatic capacity may reduce risk.

A recent position statement from professional societies lament a general disconnect between HRV as a research tool and practical clinical use (*12*). Among the barriers for clinical use, include assessments in relatively small selected cohorts, requirement for continuous ECG monitoring, and substantial variation by age, sex, and time of day.

In recent years, the widespread availability of heart rate enabled tracking devices has caused considerable interest in HRV given potential ease of availability. Commercial wrist-worn tracking devices measure heart rate intervals through photoplethysmography (PPG) at a single point of contact. PPG devices uses multiple wavelengths of light to illuminate the skin and photo-diodes to measure the reflected light, thereby inferring changes in blood volume by measuring changes in light absorption (see for e.g. (*13–15*)). While ECG-derived HRV metrics are obtained by analyzing the RR intervals between successive beats, PPG devices resolve HRV metrics through analysis of interbeat intervals (IBI) as a proxy for the RR intervals (*16–19*). PPG devices are more susceptible to motion artifacts.

Standards for HRV were set by the European Society of Cardiology, and the North American Society of Pacing and Electrophysiology (*20, 21*). HRV can be measured in many ways: in the time domain, the frequency domain, or using graphical and non-linear techniques (for an overview of HRV and metrics, see (*22*)). In this article, we present time domain, frequency domain, and graphical domain results from the largest HRV study to-date by several orders of magnitude - from *∼* 8 Million users of Fitbit devices. We demonstrate feasibility of obtaining HRV metrics from PPG at high fidelity, define diurnal distributions of common HRV metrics by age and sex, characterize the influence of aging on HRV metrics, and the relationship between physical activity on HRV metrics.

## Data and HRV metrics

Time domain metrics are computationally straightforward and do not require contiguous data. Frequency domain calculations can be computationally expensive and require the data to be contiguous and evenly sampled, but have the benefit of separating the sympathetic nervous system (fluctuations that occur on longer time scales and hence low frequencies) and the parasym-pathetic nervous system (fluctuations mostly occur on shorter time scales and hence higher frequencies) (*22*). Time windows of 5 minutes (short term) and 24 hours (long term) are commonly considered in the literature (*20, 21*). In this work, we only consider 5 minute windows, i.e a 24 hour time series would have 288 time windows. Note that 24 hour measurements of time domain metrics will be larger than the 5 minute measurements which we discuss in this work. We consider the following well known HRV metrics:

1. SDRR: The SDRR is the standard deviation of the IBI measured over a time window of 5 minutes. Let the peaks of the blood volume occur at times *T*_0_, *T*_1_, *T*_2_*, · · ·*. The IBI are the differences between successive beats, defined as:

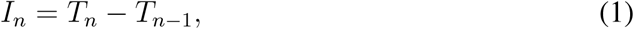

i.e. the IBI field is the first difference of the PPG waveform. The standard deviation of each 5 minute sequence of *I_n_*is computed. The SDRR measures medium to long term variations in the heart rate. SDRR correlates with the total power since the variance in the time domain equals the total power in the frequency domain. In the literature, this quantity is often termed SDNN which implies that ectopic beats are filtered out (*22*). Since we do not do this, we prefer the term SDRR.
2. RMSSD: The RMSSD is the root mean squared (RMS) value of the successive differences of the *I_n_*. The successive differences Δ*I_n_*are defined as:

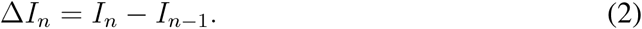 Since the Δ*I_n_* is the second difference of the PPG waveform (or the first difference of the IBI), it preferentially contains high frequency variations. The RMSSD is the RMS value of the Δ*I_n_* (i.e. the square root of the mean of the square of the samples), measured over a time window of 5 minutes. The RMSSD measures short to medium term variations in the heart rate, and correlates with HF power.
3. LF Power: The LF band measures power in the frequency range 0.04 Hz - 0.15 Hz (corresponding to physiological processes that act on timescales 6.7s - 25s), and captures both sympathetic and parasympathetic activity. Mayer waves, i.e. arterial blood pressure waves are seen in the LF band (typically around 0.1 Hz). Some vagally mediated power may also be present in the LF band, particularly during slow, paced breathing (see for example (*22*) and references therein).
4. HF Power: The HF band measures power in the frequency range 0.15 Hz - 0.4 Hz (corresponding to physiological processes that act on timescales 2.5s - 6.7s), and is a probe of the parasympathetic nervous system. The respiration induced sinus arrhythmia is usually contained in the HF band.
5. Poincare *S*_1_: The standard deviation measured along the minor axis of the Poincare ellipse is called *S*_1_, and is a measure of short term variability.
6. Poincare *S*_2_: The standard deviation measured along the major axis of the Poincare ellipse is called *S*_2_, and is a measure of long term variability.

It is well known that respiration modulates the heart rate due to the activity of the vagus nerve, at frequencies *≈* the respiration rate *∼* 10 − 20 times a minute during sleep, a phenomenon known as sinus arrhythmia (SA) (see for example (*22*) and references therein). The magnitude of the SA provides a measure of parasympathetic cardiovascular response to respiration. HRV is also a probe of sympathetic activity at lower frequencies: Mayer waves are oscillations of arterial pressure occurring spontaneously, and are enhanced during states of sympathetic activation (*23*). Fig. 1 shows the Power Spectral Density (PSD), i.e. power per unit frequency, for a single individual, measured over one night. The PSD shows two features:

1. Peak at *≈* 0.1 Hz corresponding to the arterial blood pressure induced Mayer wave.
2. Peak at *≈* 0.3 Hz corresponding to the respiration induced sinus arrhythmia.

**Figure 1:**
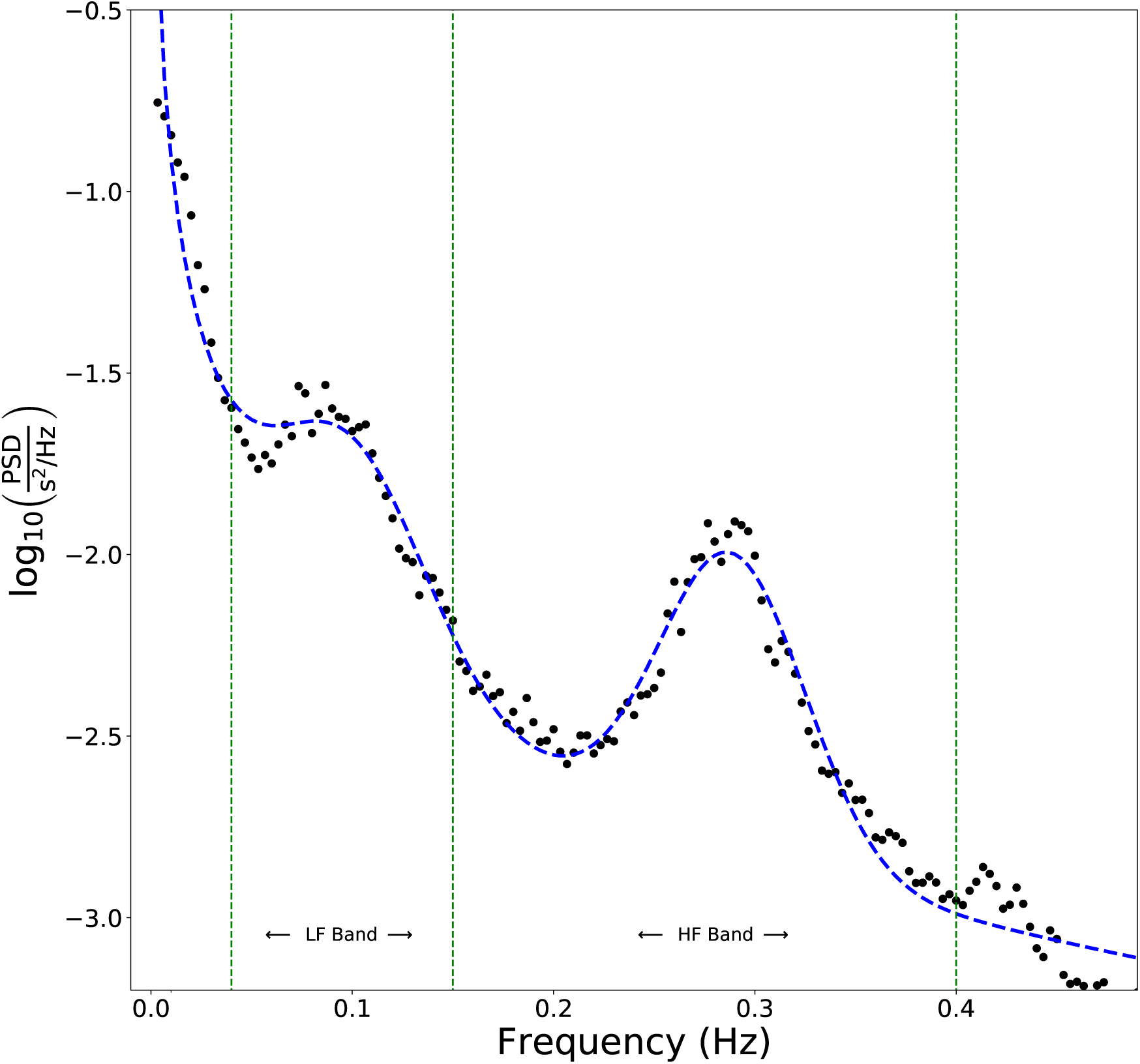
Power Spectral Density measured for a single individual over one night, showing 2 features: The Mayer wave centered at 0.0975 Hz corresponding to arterial blood pressure oscillations with a timescale of 10.3 s, and sinus arrhythmia due to respiration centered at 0.288 Hz implying a respiration rate of 17.3 breaths per minute.

We fit the data to the following form:

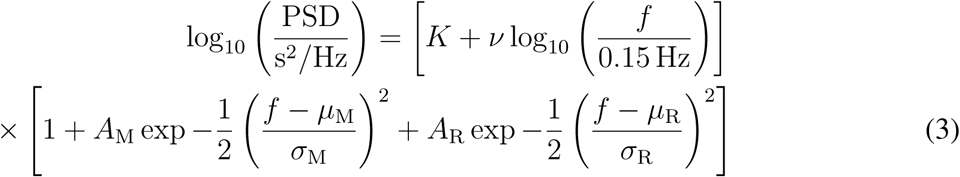

The first term represents a power law decline with frequency. The second term accounts for the two features. In this example, the best fit value for the Mayer wave is *µ*_M_ = 0.0975 Hz representing arterial pressure oscillations on a mean time scale *≈* 10.3 s. The best fit value for the SA feature for this subject was found to be *µ*_R_ = 0.2879 Hz, corresponding to a mean respiration rate *≈* 17.3 breaths per minute.

Among graphical domain techniques, we consider only first order lag-1 Poincare plots. Poincare ellipses can be categorized by their shape into a number of classes (*24–26*). Esperer et al. (*25*) investigated Poincare plots from healthy and symptomatic individuals and identified 10 distinct classes, each with diagnostic value. Some classes of Poincare ellipses obtained from our data, are shown in Fig. 2. The top row shows heart beats in sinus rhythm. (a) is the “comet class” and represents a healthy heart. (b) is termed the “torpedo” class since it lacks the taper seen in the comet plots. Tulppo et al. (*26*) distinguish the comet and torpedo classes based on the ratio of short and long axes. Esperer et al. (*25*) instead, distinguish these classes based on the HRV. The torpedo shaped Poincare plots show cardiovascular dysfunction since the short term variability is weaker than what is seen in the comet plots. (c) is an example of tachycardia, and shows very low variability with substantially reduced *S*_1_ and *S*_2_. The bottom row shows examples of arrhythmias. (d) shows the “fan” pattern suggestive of atrial fibrillation (*24, 25*), while (e) and (f) show bi-lobed and tri-lobed anomalies which could potentially indicate premature atrial or ventricular beats.

**Figure 2:**
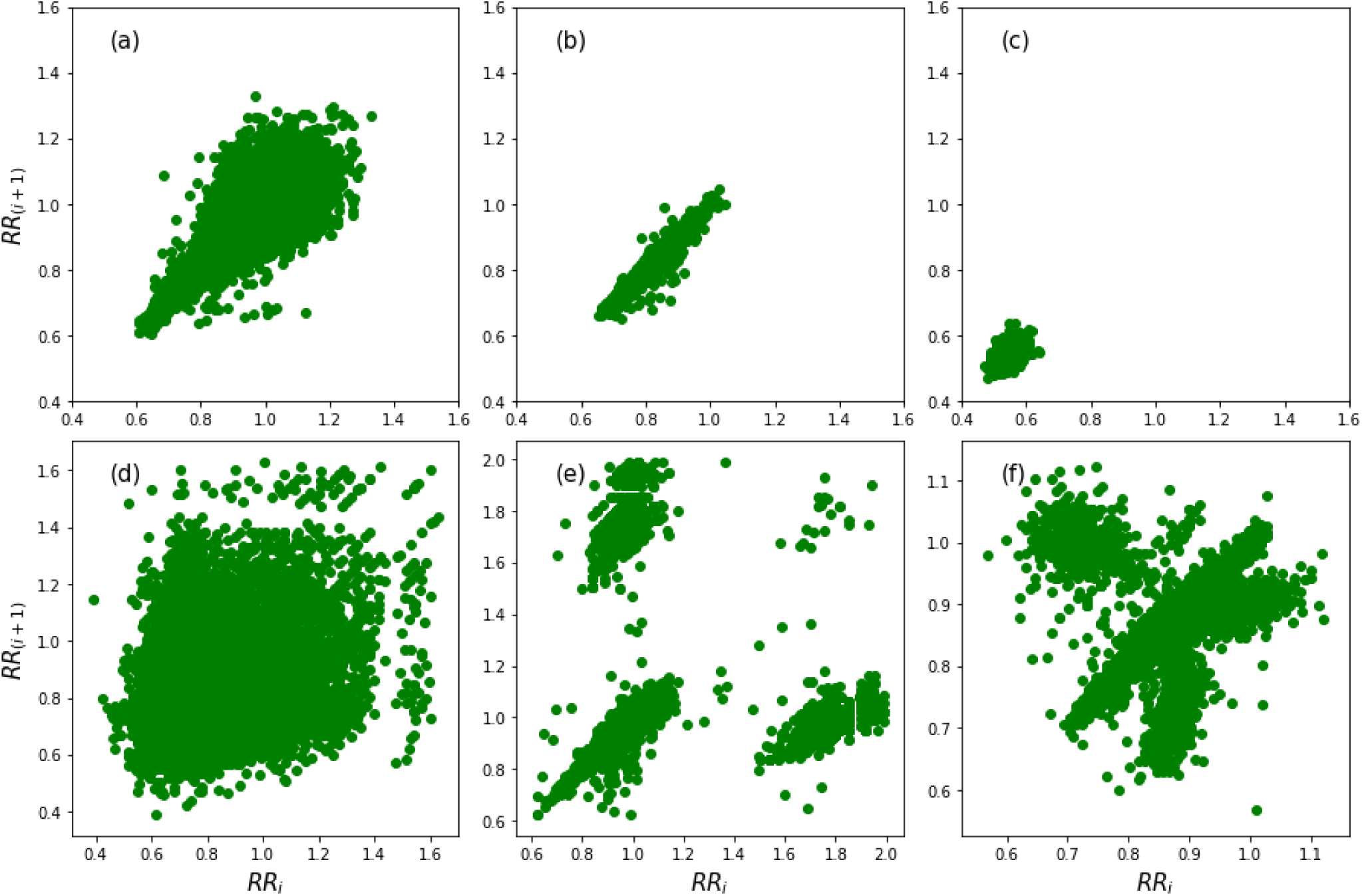
Poincare plots are very useful in diagnosing heart arrhythmias. (a) shows the “comet” pattern of a healthy heart. (b) shows the “torpedo” pattern indicating low short term variability. (c) is an example of a heart in tachycardia. (d) shows the fan pattern suggestive of atrial fibrillation. (e) and (f) show bi-lobed and tri-lobed patterns which could indicate premature atrial and ventricular contractions.

### Changes in HRV with age

It is well known that HRV declines with age, although the decline may be ameliorated by healthy habits, e.g. staying active, mindfulness practices, etc. Umetani et al. (*1*) studied the decline in time domain HRV metrics with age, from a dataset of 240 healthy subjects. They found that the RMSSD declines more rapidly than the SDRR (called SDNN Index in their paper). They also found that for age *<* 30 yr, HRV in female subjects was lower on average than in male subjects for all time domain metrics, with no gender differences for age *>* 50 yr. Other works that discuss HRV and aging include (*27–29*).

Fig. 3 shows the decline in HRV with aging (from age 20 yr to 60 yr), from our data. Our results are consistent with the findings of Umetani et al. (*1*) especially for measurements taken during the daytime (our SDRR corresponds to their SDNN Index). Interestingly, the decline in HRV with age depends not only on gender, but also the time of day when the measurements are made. Plots (a) and (b) show the decrease in RMSSD and SDRR respectively. The SDRR is higher on average, in men compared to women, a trend that is less noticeable with the RMSSD. Plots (c) and (d) show equivalent results for the HF power and LF power, while plots (e) and (f) show the variation in Poincare *S*_1_ and *S*_2_. Note that the HF power, Poincare *S*_1_, and RMSSD behave similarly. Similarly LF power, Poincare *S*_2_, and SDRR show a high degree of correlation. The RMSSD tends to decline faster than the SDRR, the HF power declines faster than the LF power, and the Poincare *S*_1_ declines faster than *S*_2_. This suggests that with increasing age, parasympathetic ability is lost sooner than sympathetic ability.

### Changes in HRV with time of day

Vandewalle et al. (*30*) studied the diurnal variation of Heart Rate Variability metrics over a 24 hour period involving eight healthy male subjects, and found that HRV metrics vary throughout the day, reaching peak values in the early morning hours. Our data also shows that HRV metrics vary significantly throughout the day, and hence HRV measurements should be taken consistently at the same time of day. Fig. 4 shows the diurnal variation of HRV metrics. Plots (a) and (b) show the fractional variation of the RMSSD and the SDRR as a function of the time of day, for male (green) and female (red) subjects. Young users (age = 20-21 yr) are represented by solid lines, while older users (age = 60-61 yr) are represented by dashed lines. Plots (c) and (d) show the variation for HF power and LF power. Similarly, plots (e) and (f) show the daily modulation in *S*_1_ and *S*_2_. The SDRR, LF power, and Poincare *S*_2_ show a change in phase with increase in age: Older users tend to have an earlier peak in the daily cycle, for the sympathetic measures. All HRV metrics peak early in the day (*∼* between 5 am and 8 am) and reach a minimum in the late evening (*∼* 7 pm - 8 pm).

**Figure 3:**
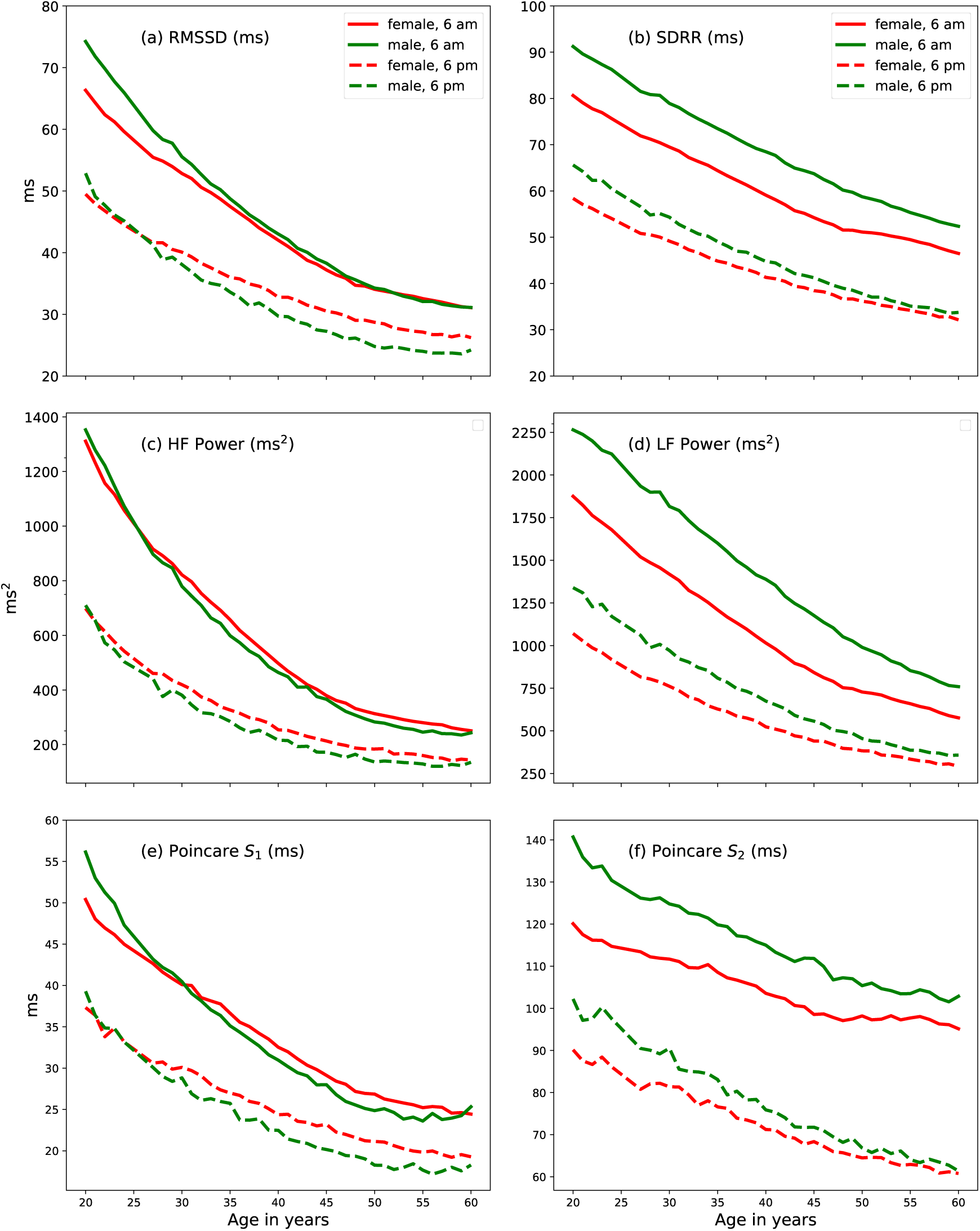
Decline in HRV with age. All metrics decrease with age, for both men and women. However, the parasympathetic measures (RMSSD, HF power, and Poincare *S*_1_) decline faster than the sympathetic measures (SDRR, LF power, and Poincare *S*_2_).

**Figure 4:**
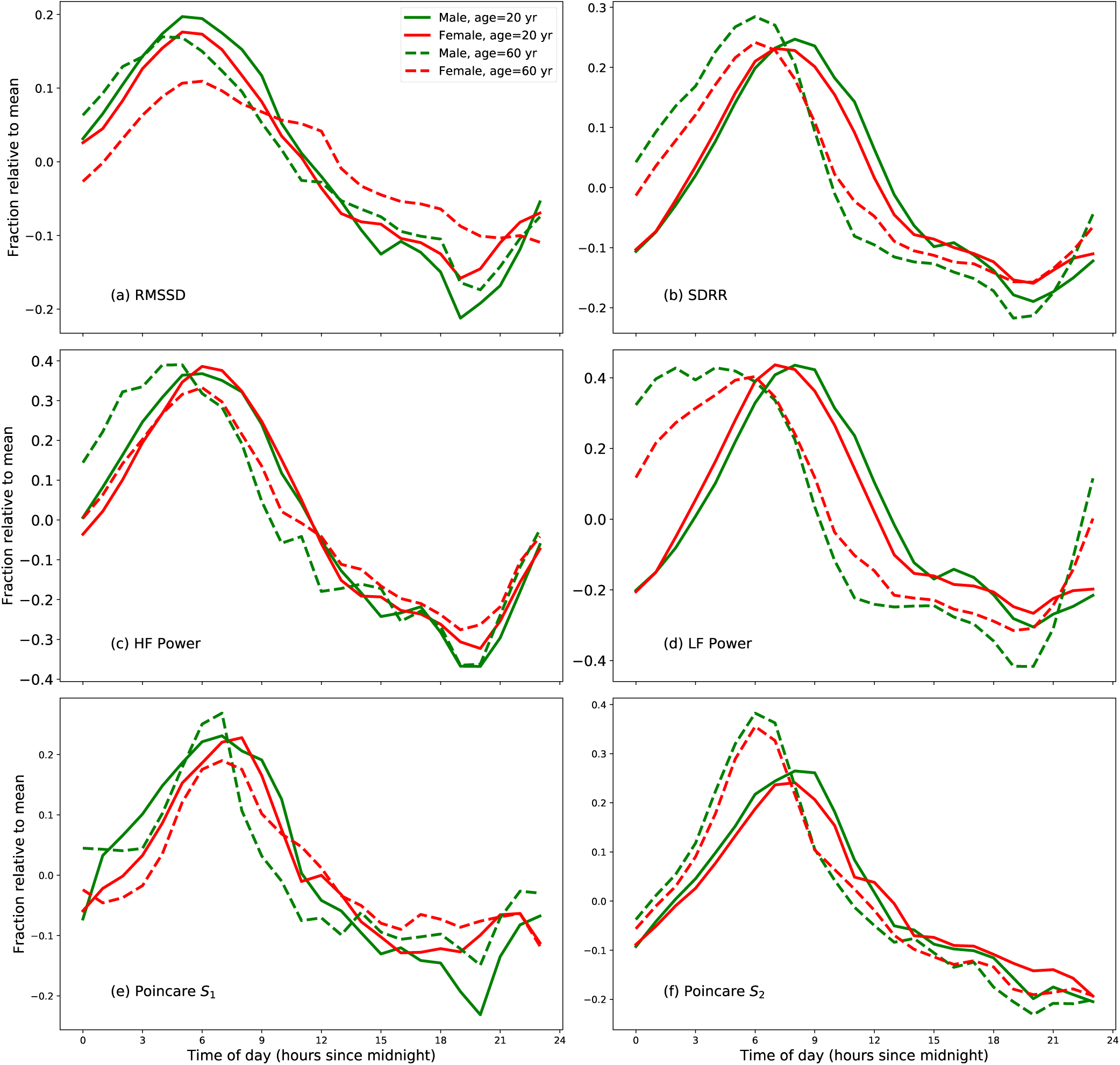
Daily variation of HRV features. The modulation is substantial for all ages. The phase of the sympathetic measures (i.e. SDRR, LF power, and Poincare *S*_2_) shifts to earlier times for older individuals. The change in phase with age is less prominent in the modulation of the parasympathetic measures (RMSSD, LF power, and Poincare *S*_1_).

### Scaling relations

In this section, we take a closer look at the scaling of HRV with age, gender, and time of day. We parametrize the HRV by the following power law form:

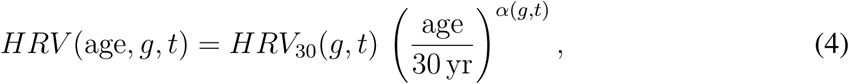

where *t* is the time of day in hours from midnight, ‘age’ is the age in years, and *g* is the gender. *HRV* is the HRV metric being studied, which could be HF power, LF power, RMSSD, SDRR, *S*_1_, or *S*_2_. Note that the dependence on the time of day and the age are not separable, i.e. the power law exponent *α* is a function of the time of day.

The time dependance of *α* means that the decline in HRV with age is different at different times of the day. Let us expand the time dependent terms in Eq. 4, i.e. *HRV*_30_ and *α* as follows:

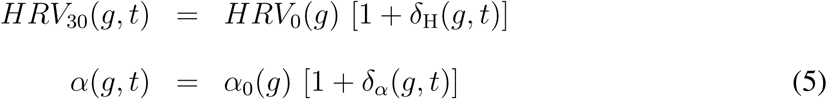

*HRV*_0_(*g*) and *α*_0_(*g*) are mean values (over 24 hours) and are gender dependent. Table 1 gives the values of *α*_0_(*g*) and *HRV*_0_(*g*) for the different HRV metrics. Together with the time dependent terms, one can compute the HRV given the gender, age, and time of day. Fig. 5 shows the diurnal modulation of *δ*_H_(*g, t*) and *δ_α_*(*g, t*). The plots for *δ*_H_(*g, t*) are sufficiently close to sinusoidal, that we can approximate *δ*_H_(*g, t*) using the first 3 Fourier components:

**Figure 5:**
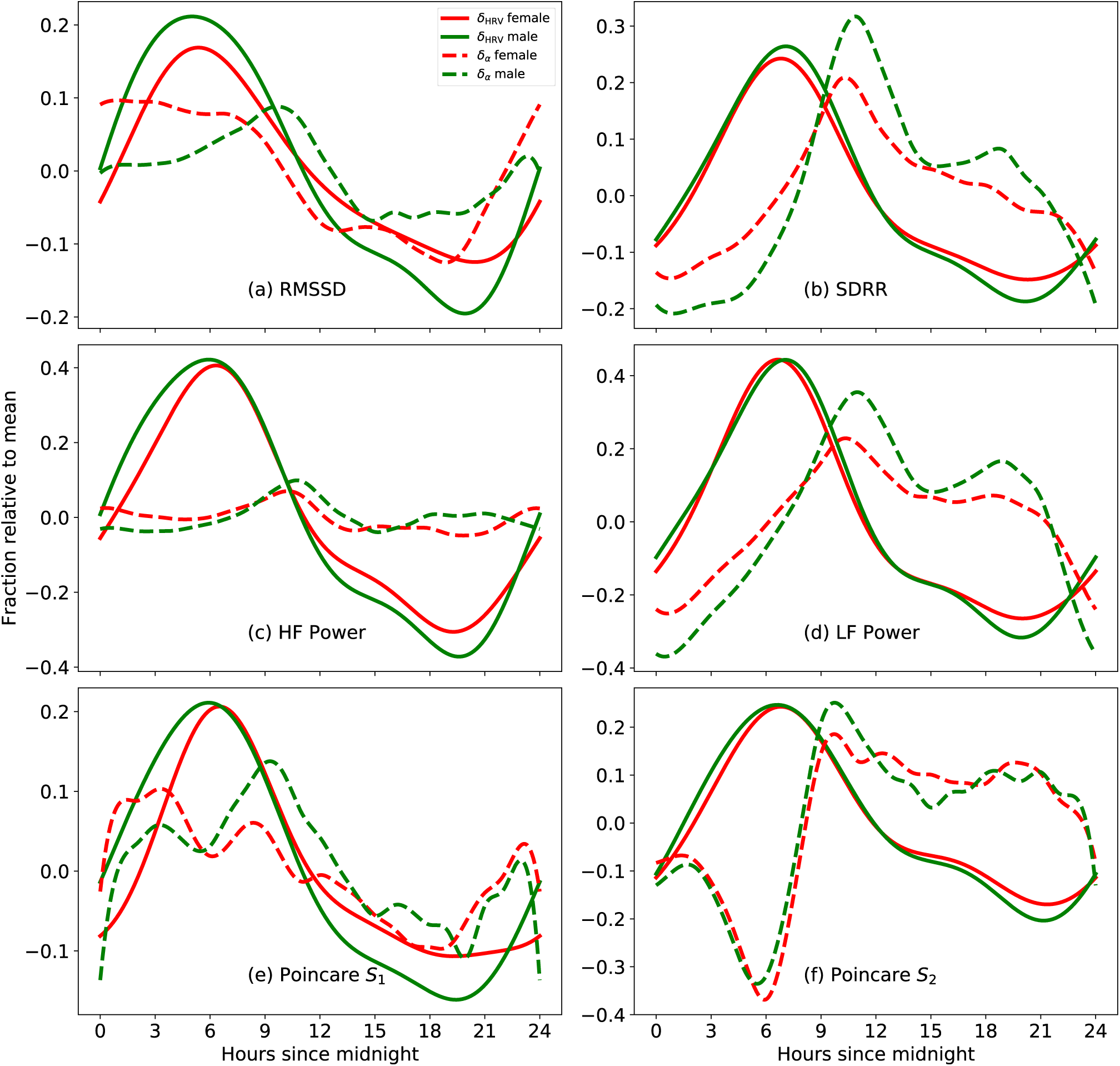
Daily variation of the time dependent scaling parameters: *δ*_H_ and *δ_α_*. The variation of *δ_α_* is larger for the sympathetic measures (SDRR, LF power, Poincare *S*_2_) compared to the parasympathetic measures (RMSSD, HF power, Poincare *S*_1_). This helps explain the change in phase seen in Fig. 4

**Table 1:**
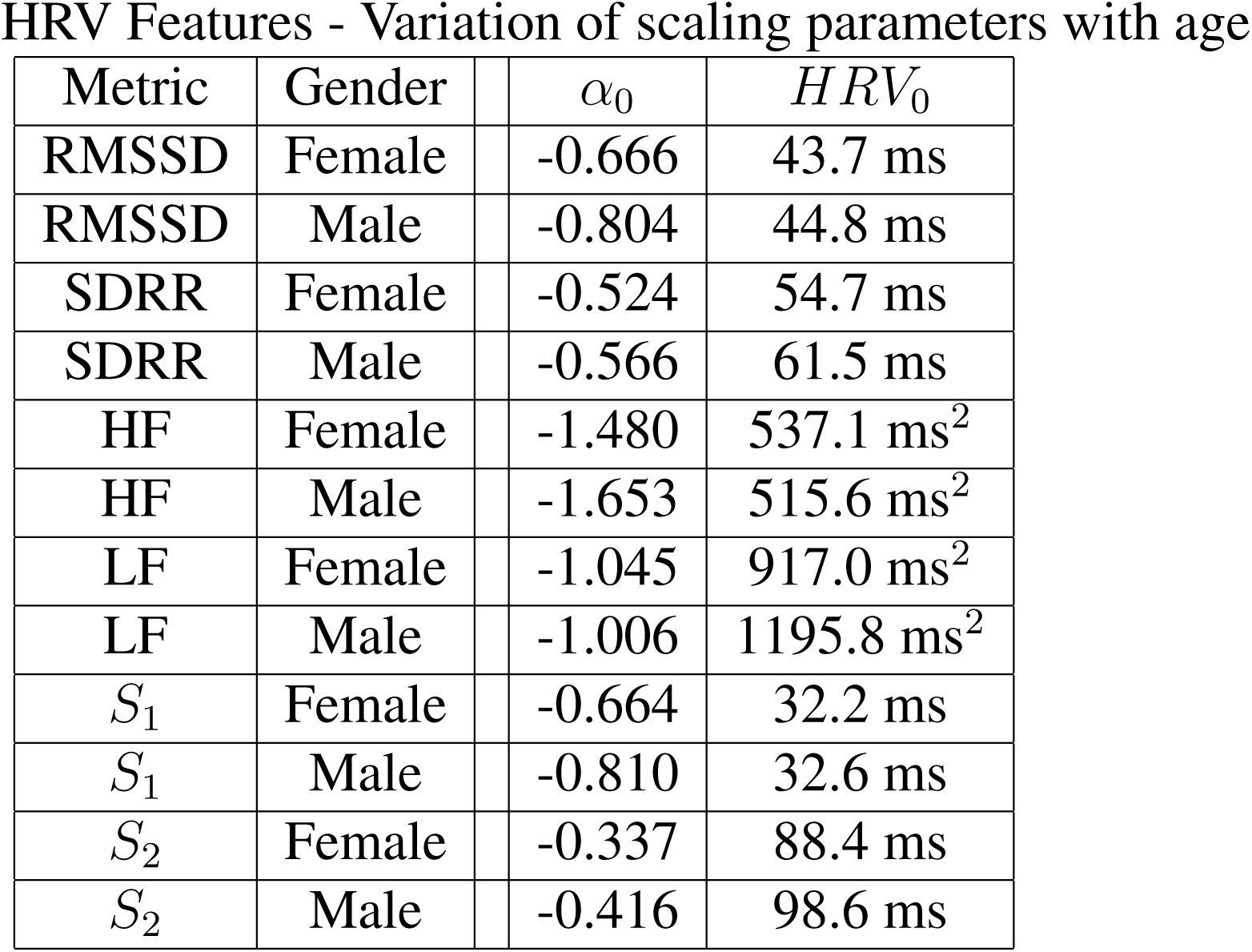
The scaling parameters *α*_0_(*g*) and *HRV*_0_(*g*)

| Metric | Gender | $\alpha_0$ | $HRV_0$ |
| --- | --- | --- | --- |
| RMSSD | Female | -0.666 | 43.7 ms |
| RMSSD | Male | -0.804 | 44.8 ms |
| SDRR | Female | -0.524 | 54.7 ms |
| SDRR | Male | -0.566 | 61.5 ms |
| HF | Female | -1.480 | 537.1 ms<sup>2</sup> |
| HF | Male | -1.653 | 515.6 ms<sup>2</sup> |
| LF | Female | -1.045 | 917.0 ms<sup>2</sup> |
| LF | Male | -1.006 | 1195.8 ms<sup>2</sup> |
| $S_1$ | Female | -0.664 | 32.2 ms |
| $S_1$ | Male | -0.810 | 32.6 ms |
| $S_2$ | Female | -0.337 | 88.4 ms |
| $S_2$ | Male | -0.416 | 98.6 ms |

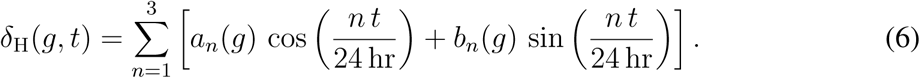

The values of *a_n_*(*g*) and *b_n_*(*g*) for *n* = 1,2,3 are tabulated in Table 2, for male and female subjects. In all cases, *b*_1_ is the dominant term, describing a sinusoid with a period of 24 hours. The other terms are corrections to the sinusoidal variation. Tables 1 and 2 give the value of typical HRV metrics for a 30 yr old person. More accurate estimates can be obtained by accounting for the time dependent term *δ_α_*(*t*) as shown in Fig. 5. Typically HRV numbers for other ages can be obtained from Eq. 4, and from Tables 1 and 2.

**Table 2:** Variation of time dependent scaling parameters: The first 3 Fourier components.

| Metric | Gender | $a_1$ | $a_2$ | $a_3$ | $b_1$ | $b_2$ | $b_3$ |
| --- | --- | --- | --- | --- | --- | --- | --- |
| RMSSD | Female | -0.009 | -0.029 | -0.003 | 0.137 | 0.022 | 0.001 |
| RMSSD | Male | 0.012 | -0.02 | 0.011 | 0.196 | 0.035 | 0.011 |
| SDRR | Female | -0.047 | -0.051 | 0.01 | 0.173 | -0.012 | -0.007 |
| SDRR | Male | -0.052 | -0.044 | 0.019 | 0.2 | -0.01 | -0.001 |
| HF | Female | -0.02 | -0.06 | 0.025 | 0.323 | 0.015 | -0.02 |
| HF | Male | 0.021 | -0.046 | 0.033 | 0.382 | 0.044 | 0.005 |
| LF | Female | -0.06 | -0.098 | 0.022 | 0.317 | -0.014 | -0.016 |
| LF | Male | -0.067 | -0.073 | 0.043 | 0.335 | -0.015 | -0.004 |
| S1 | Female | -0.035 | -0.05 | 0.003 | 0.137 | -0.010 | -0.015 |
| S1 | Male | 0.007 | -0.03 | 0.009 | 0.181 | 0.0130 | 0.0 |
| S2 | Female | -0.063 | -0.061 | 0.010 | 0.170 | -0.002 | -0.002 |
| S2 | Male | -0.059 | -0.056 | 0.009 | 0.193 | 0.014 | 0.007 |

Focusing on Fig. 5, plot (a) show the daily modulation of *δ*_RMSSD_ (solid lines) and *δ_α,_*_RMSSD_ (dashed lines) for male (green) and female (red) subjects. Plot (b) shows the equivalent curves for the SDRR, while plots (c), (d), (e), and (f) show *δ*_H_ and *δ_α_*for the HF power, LF power, *S*_1_ and *S*_2_. The variation of *δ*_H_ is significant in all cases. The variation of *δ_α_* is small for HF power, significant for the RMSSD and Poincare *S*_1_, and large for SDRR, LF power, and Poincare *S*_2_. In light of this figure, we can better understand the phase variation seen in Fig. 4 which showed that the phase of the SDRR, LF, and *S*_2_ modulation peaks earlier in older individuals. From Fig. 5, we note that *δ_α_* changes rapidly when *δ*_H_ is close to a maximum. This means that as people get older, the SDRR/LF/*S*_2_ at *∼* 7 am declines faster than the same metrics, at e.g. *∼* 6 am, explaining the movement of the phase with increase in age. Such as effect is not apparent in the RMSSD, HF power, or Poincare *S*_1_ because *δ_α_* varies slowly when *δ*_H_ is a maximum.

Finally, we investigate the possibility of increasing HRV through behavior modification. Several authors (*31–34*) have discussed the effect of physical activity on HRV and studies have shown beneficial results. We analyzed the HRV of all participants (measured from 6 am - 7 am) grouped by the average number of steps taken per day (steps per day is averaged over a 90 day period preceding the HRV measurement). Fig. 6 shows a strong correlation between physical activity and HF and LF power (similar conclusions hold true for other metrics). Plots (a) and (b) show the change in HF and LF power respectively for younger users (ages 20-24 yr). Corresponding results for older users (ages 50-54 yr) are shown in plots (c) and (d). To estimate the impact of exercise, we modeled the HRV power variation by a linear fit: HRV power = *C* + steps*/σ*, where *σ* is the number of steps necessary to increase the power by 1 ms^2^ on average. The values of Pearson correlation coefficient (*r*), *C*, and *σ* are listed in Table 3. An alternative approach to increasing HRV involves the practice of mindfulness or meditation techniques (*35–37*).

**Figure 6:**
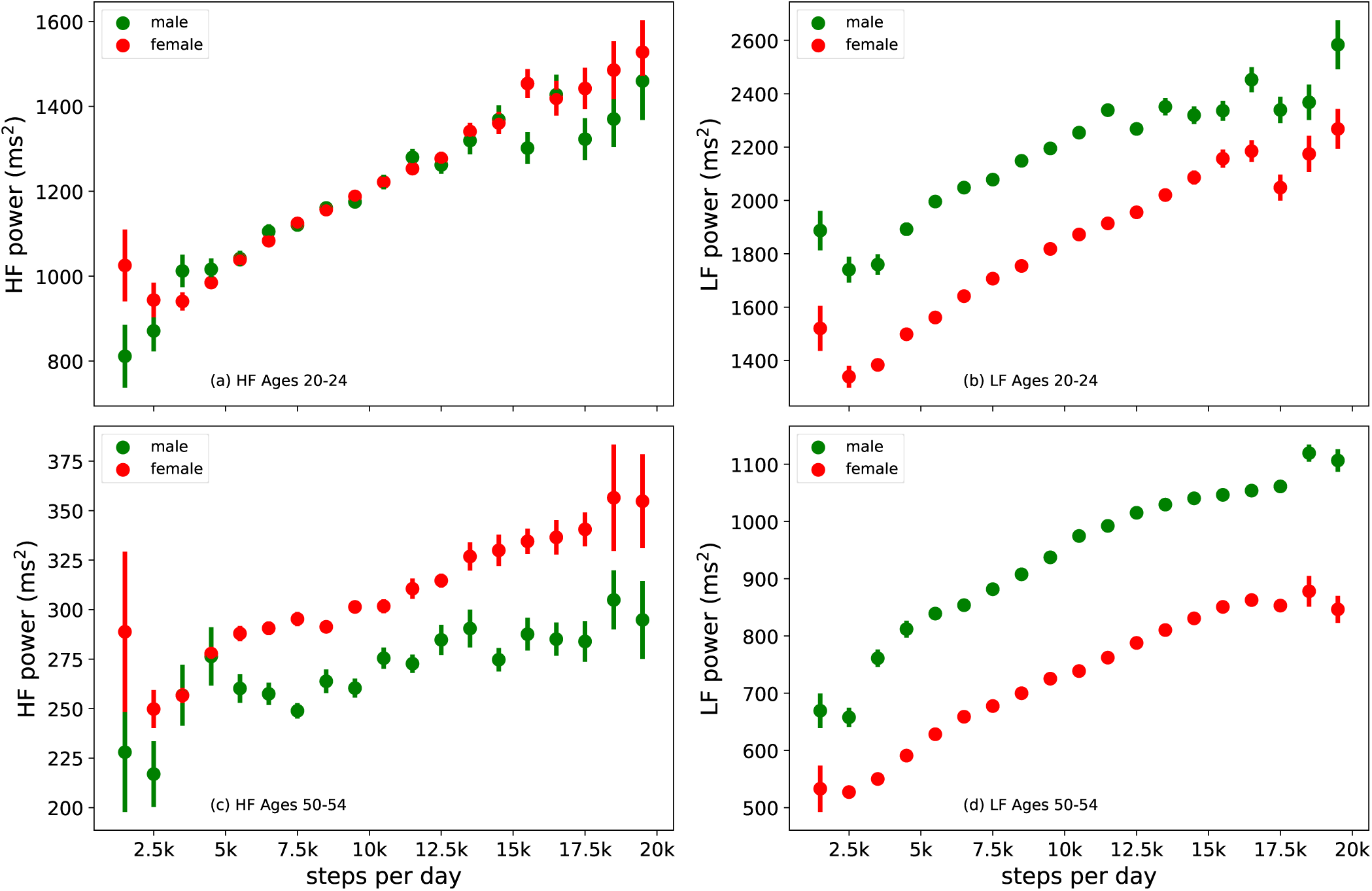
HRV can be increased with physical exercise for all ages, but especially for younger individuals. Shown are HF and LF variations (measured from 6 am - 7am) with the number of steps per day, for young (ages 20-24 yr) and older (ages 50-54 yr) participants.

**Table 3:** Effect of activity on HRV power. People of all ages can improve their HRV through physical activity, but the effect is larger for younger users, especially for HF power.

| Gender | Metric | Age (yr) | $r$ | C (ms<sup>2</sup>) | $\sigma$ (steps) |
| --- | --- | --- | --- | --- | --- |
| Male | HF | 20-24 | 0.961 | 865 | 32 |
| Female | HF | 20-24 | 0.985 | 873 | 30 |
| Male | LF | 20-24 | 0.946 | 1758 | 25 |
| Female | LF | 20-24 | 0.973 | 1314 | 20 |
| Male | HF | 50-54 | 0.859 | 234 | 295 |
| Female | HF | 50-54 | 0.954 | 254 | 195 |
| Male | LF | 50-54 | 0.969 | 683 | 42 |
| Female | LF | 50-54 | 0.979 | 510 | 48 |

## Discussion

In this article, we presented heart rate variability measurements from 8 Million individuals using Fitbit devices, taken over a period of 24 hours. We reported results on HRV metrics in the time domain, frequency domain, and the graphical domain. To compute frequency domain metrics, we first interpolated the IBI field to obtain samples that are evenly separated. The PSD was computed from the interpolated field, through a fast fourier transform. The spectral shape of the PSD of a healthy subject contains two features in addition to a power law: Respiration induced sinus arrhythmia at *∼* 0.2 − 0.3 Hz, and the arterial blood pressure induced Mayer wave around *∼* 0.1 Hz. We showed examples of Poincare plots for subjects with heart beats in sinus rhythm, as well as for subjects with arrhythmias.

We have provided benchmark tables for HRV features in the Supplementary text giving the mean, median and the 25^th^− 75^th^ percentile ranges, for different ages, and for both male and female participants. We anticipate that a user’s HRV in relation to the HRV of others of similar age and gender may be a useful feature in assessing cardiovascular risk. Popular risk assessment algorithms such as the Framingham risk score, do not take HRV into account and sometimes underestimate the risk of adverse cardiovascular events (*38*) and sometimes overestimate it (*39*). Authors (*40*) found that the Framingham risk score for men was inversely correlated with several HRV features. We are therefore optimistic that a person’s HRV relative to others in their gender and age group would help improve cardiovascular risk predictions.

Our results may have important implications for the remote monitoring of human health given the widespread availability of wrist-worn trackers. First, our method allows for the continuous monitoring of autonomic and cardiovascular responses throughout life’s experiences. Consistent with prior small studies, all HRV metrics decrease with age (*1,41,42*). The RMSSD, HF power, and Poincare *S*_1_ decrease with age faster than the SDRR, LF power, and Poincare *S*_2_. This suggests a more rapid decline of parasympathetic function with increasing age, compared to sympathetic activity. The LF/HF ratio shows an *increase* with age up to *≈* 50 − 60 yr, also implying a faster decline in parasympathetic function. Prior work in older adults implies the the slope of decline in parasympathetic HRV metrics is inversely related to longevity (*42*). We also presented results showing the diurnal variation of HRV metrics. The variation is substantial, and hence it is advisable for people to interpret HRV measurements at the same time of day. With the help of this massive dataset, we have been able to show a difference between sympathetic and parasympathetic measures with respect to the phase variation of the daily modulation with age. The sympathetic measures show a phase shift towards earlier times of the day with increase in age. Such an effect is far less noticeable with the parasympathetic measures.

Second, we presented simple formulae and tables that show how to compute typical HRV metrics from photoplethysmography for interpretation. We fitted the variation of HRV metrics with age, using a simple power law. The scaling parameters depend on gender, and time of day. The HRV metrics can be estimated given age, gender, and time of day using 2 gender dependent scaling parameters *HRV*_0_(*g*), *α*_0_(*g*), and 2 gender and time dependent scaling parameters *δ*_H_(*g, t*), and *δ_α_*(*g, t*). The parameter *δ*_H_(*g, t*) can be well approximated using the first 3 Fourier components. The time variation of *δ_α_*(*g, t*) is more complicated, and we present figures that allow for quick estimations.

Third, since we observe a strong correlation between physical steps and HRV, increasing physical activity may optimize HRV metrics. While there have been other studies of exercise and HRV, they have been typically limited to small populations (*31, 32*) or to participants in a narrow age range (*33*). Due to the size of our dataset, we have been able to examine the correlation between exercise and HRV in more detail, for both young and older participants. To estimate the increase in HRV due to physical activity, we modeled the variation of HRV power with activity, by a linear approximation. The inverse slope of this curve gives us the number of additional steps per day necessary to yield a 1 ms^2^ increase in power. Users of all ages may optimize their HRV through physical exercise, although the effect is larger for younger subjects, especially for HF power. The linear fit models suggest that people in the age range 20-24 yr may increase their HF power by 1 ms^2^ with every *∼* 30 additional steps. By contrast, subjects in the age range 50-54 yr need *∼* 200 (female) - 300 (male) additional steps for each 1 ms^2^ increase in HF power. As a result, older individuals improve their LF power more than HF power, with physical activity. Such a large difference in HF recovery with age suggests that parasympathetic function is harder to restore with physical activity, and is an important finding of the present work. While HRV metrics have been previously correlated with cardiovascular health and mortality, our technical advance and descriptions now permit its potential use for health promotion through tens of millions of currently available wrist-worn commercial trackers. Randomized controlled trials are now necessary to demonstrate effective ways to use HRV metrics to improve health.

## Acknowledgments

We thank Emily Blanchard, Eric Chang, Robert da Silva, Tony Faranesh, Karla Gleichauf, Suraj Gowda, Sarah Kernasovskiy, Belen Lafon, Lindsey Sunden, Teresa Tenfelder, and Shelten Yuen for many helpful discussions. A.N., A.P., and H. E.-F. are employees of Fitbit Inc. P.N. is supported by awards from NHLBI (R01HL142711, R01HL148050, R01HL148565) and a Hassenfeld Scholar Award from the Massachusetts General Hospital. P.N. also reports grants from Amgen, Apple, and Boston Scientific, and consulting income from Apple and Blackstone Life Sciences, all unrelated to the current work.

## Supplementary materials

### Data and Methods

The output of a PPG device is the Interbeat Interval (IBI) tachogram, i.e. the time between peaks of blood volume. The IBI are susceptible to noise due to motion artifacts, electronic noise, missed heart beats, etc. It is essential to clean the IBI field if it is to be a faithful representation of the RR interval tachogram measured by an electrocardiogram (ECG). Cleaning the noisy PPG data is accomplished by a proprietary technique consisting of a median filtering stage, and an anomaly detection stage. Fig. S1 shows the comparison between PPG and ECG, for 60 minutes of data. PPG (with fitbit devices) and ECG (single lead, chest strap) data were simultaneously obtained from *∼* 30 subjects (normal sinus rhythm, users were at rest, data was collected with IRB approval). The data was then interpolated so that PPG and ECG readings could be compared at the same instant of time. Fig. S1(a) shows the correlation between the interpolated RR intervals collected by the ECG device, and the interpolated IBI (henceforth IIBI) obtained from PPG. The red points represent raw IIBI, whereas the green dots represent the cleaned IIBI. Fig. S1(b) shows the total power (i.e. power spectrum of the IBI time series data integrated over all frequencies) contained in the data (the power is computed over 5 minute windows). The red points are total power calculations from the raw IIBI, whereas the green points represent total power computed from the cleaned IIBI. For the comparison of total power (plot (b)), the Pearson correlation coefficient between the ECG derived data and the cleaned PPG data is *r* = 0.96, implying a high degree of correlation between PPG derived HRV metrics and ECG derived HRV metrics. In contrast, the correlation is only *r* = 0.39 for the raw data.

It is clear from Fig. S1(b) that cleaning the PPG data is necessary before computing HRV features. Our cleaning algorithm does not make an assumption about noise levels. We define noise by the prominence, i.e. the value in relation to its neighboring points. Since very noisy samples can affect the prominence, we choose to discard these samples. We discard the data in a time window if the fraction of noise spikes exceeds *∼* 10% of the window size. For the remainder of the paper, we will describe HRV metrics derived from the cleaned PPG data.

We collected data from *∼* 8 Million individuals using Fitbit devices over the course of 24 hours. The data was anonymized, and the analysis was consistent with Fitbit’s terms and conditions. Table S1 describes the number of individuals in our study, by age and gender. The columns labeled “Fem. (any)” and “Male (any)” are the number of female and male participants with HRV data at some time of the day. The columns labeled (6 am) and (6 pm) show the number of participants with data at 6 am - 7 am and at 6 pm - 7 pm respectively. These times of day are close to the maxima and minima of the HRV circadian rhythm, although these vary by age and gender. We were able to obtain significantly more usable data when the subjects were asleep due to the absence of motion artifacts. Data coverage at 6 am - 7 am for female (male) participants decreases from 70% (65%) for younger users, to 61% (56%) for older users. The coverage at 6 pm - 7 pm for female (male) participants increases from 7.4% (6.6%) to 14% (14%) for older users.

The distribution of HRV metrics is shown in Fig. S2, for subjects of age 30 - 31 yr, for male (green bars) and for female (red bars) participants. The HF power and LF power are strongly skewed, with the mean larger than the median. The RMSSD and Poincare *S*_1_ are also skewed, while the SDRR and Poincare *S*_2_ show a smaller skew. Men have a larger SDRR, LF power, and Poincare *S*_2_ on average compared to women. The difference in gender is less apparent in the RMSSD, HF power, and the Poincare *S*_1_. To estimate the difference in parasympathetic and sympathetic HRV measures between male and female participants, we performed a *t*-test to quantify the difference as a function of age and time of day. When measured at 6 am - 7 am, the HF power is higher in female subjects for ages *>* 33 yr, while male subjects have a higher HF power for smaller ages. When measured at 6 pm - 7 pm, the HF power is higher in female participants for ages *>* 24 yr. We find that male participants always have higher LF power compared to female participants, but the difference is larger in the morning compared to the evening.

We compute the standard deviation (SDRR) of the IBI, and the root mean squared value of the successive differences (RMSSD), over time windows of size 5 minutes. Time domain HRV metrics are sensitive to the size of the input window since Fourier modes longer than *W* are not contained in a time window of size *W* .

Frequency domain calculations require further pre-processing: we resample the IBI field to obtain 512 equally spaced samples in each 5 minute time window. We can thus resolve all frequency components up to 0.5 *×* (512*/*300) *≈* 0.85 Hz. We do not expect significant power at frequencies above 0.5 Hz. The resolution in frequency space is 1/300 Hz which gives us an adequate number of samples in the LF and HF bands. The mean of the data in the time window is subtracted, and the IIBI field is smoothed with a Hann window. A Fast Fourier Transform is then applied to the smoothed IIBI field, and properly normalized to give us the Power Spectral Density (PSD), which is the power contained in the IIBI field per unit frequency. Finally, the PSD is integrated over the relevant frequencies to give us the band power. Frequency domain analysis requires the data in a window to be contiguous and evenly sampled. Therefore, missing data will need to be imputed through interpolation. As a result, we do not consider data windows with a coverage lower than 70%.

Poincare plots are scatter plots obtained by plotting the IBI at time index *i* against the succeeding IBI, i.e. at time index *i* + 1 (more accurately, these are called first order lag-1 Poincare plots). The Poincare plots will contain a high density of points scattered close to the 45*^◦^* line. This scatter is a measure of variability. Fig. S3 shows the Poincare plot for a healthy subject. The plot resembles a tapered ellipse since larger IBI allow for more variability. The standard deviation of points along the major axis (*S*_2_) is a measure of long term variability, while the standard deviation along the minor axis (*S*_1_) is a measure of short term variability.

Table S2, Table S3, and Table S4 show benchmark values for time domain, frequency domain, and graphical domain HRV metrics respectively. The HRV metrics are shown for various ages, for female and male users, and at two times of the day (from 6 am - 7 am, and from 6 pm - 7 pm). We provide the mean, the median, as well as the range from the 25*^th^* percentile to the 75*^th^* percentile. This range is computed over our entire population for a specific age, gender, and time of day. The measurement from 6 am - 7 am indicates a median over all available 5 minute windows. For a window to be acceptable, the coverage should be adequate, and the noise fraction should be sufficiently small. There are a maximum of 12 such windows in an hour, and we discard the entire hour if there are less than 3 acceptable windows. Table S5 shows the LF/HF ratio which is a measure of sympathovagal balance.

**Table S1:** Number of participants in the present study, by age and gender. The amount of data depends on the time of day since we lose data during the daytime due to motion artifacts.

| Age | Fem. | Fem. | Fem. | Male | Male | Male |
| --- | --- | --- | --- | --- | --- | --- |
| (yr) | (any) | (6 am) | (6 pm) | (any) | (6 am) | (6 pm) |
| < 20 | 168k | 118k | 12k | 69k | 45k | 5k |
| 20-30 | 895k | 598k | 84k | 340k | 217k | 32k |
| 30-40 | 1.23M | 811k | 114k | 582k | 360k | 54k |
| 40-50 | 1.16M | 748k | 131k | 593k | 359k | 65k |
| 50-60 | 1.06M | 651k | 134k | 570k | 333k | 72k |
| 60-70 | 688k | 417k | 96k | 406k | 231k | 56k |
| > 70 | 248k | 150k | 36k | 188k | 105k | 27k |

**Table S2:** Typical values for time domain HRV features: The RMSSD distribution is more skewed than the SDRR distribution (see Fig. S2) and this is reflected in the difference between the mean and the median. Bin size for the age is 1 year, i.e. age = 20 includes users between 20 and 21 years of age.

HRV Features - Time Domain
| Time of day | Age<br>(yr) | RMSSD (female) |  |  | RMSSD (male) |  |  | SDRR (female) |  |  | SDRR (male) |  |  |
| --- | --- | --- | --- | --- | --- | --- | --- | --- | --- | --- | --- | --- | --- |
|  |  | (mean) | (med) | (25p-75p) | (mean) | (med) | (25p-75p) | (mean) | (med) | (25p-75p) | (mean) | (med) | (25p-75p) |
| 6am - 7am | 20 | 66 | 56 | 37 - 85 | 74 | 66 | 45 - 96 | 81 | 76 | 56 - 101 | 91 | 88 | 66 - 114 |
| 6am - 7am | 25 | 57 | 48 | 32 - 73 | 61 | 54 | 35 - 79 | 73 | 68 | 50 - 91 | 82 | 79 | 57 - 103 |
| 6am - 7am | 30 | 53 | 45 | 31 - 67 | 56 | 49 | 34 - 71 | 69 | 65 | 49 - 86 | 79 | 75 | 57 - 98 |
| 6am - 7am | 35 | 47 | 41 | 29 - 60 | 49 | 43 | 31 - 62 | 64 | 61 | 46 - 79 | 73 | 70 | 54 - 91 |
| 6am - 7am | 40 | 42 | 37 | 26 - 52 | 43 | 38 | 27 - 54 | 59 | 56 | 43 - 73 | 68 | 65 | 50 - 85 |
| 6am - 7am | 45 | 37 | 33 | 24 - 46 | 38 | 34 | 25 - 48 | 54 | 51 | 40 - 66 | 64 | 61 | 47 - 78 |
| 6am - 7am | 50 | 34 | 31 | 22 - 42 | 34 | 31 | 23 - 42 | 51 | 49 | 38 - 63 | 59 | 56 | 42 - 72 |
| 6am - 7am | 55 | 33 | 29 | 22 - 40 | 32 | 29 | 21 - 39 | 49 | 47 | 36 - 61 | 55 | 52 | 40 - 68 |
| 6am - 7am | 60 | 31 | 28 | 21 - 38 | 31 | 27 | 20 - 37 | 46 | 44 | 34 - 57 | 52 | 49 | 37 - 64 |
| 6pm - 7pm | 20 | 49 | 41 | 28 - 62 | 53 | 45 | 30 - 66 | 58 | 54 | 40 - 73 | 66 | 63 | 46 - 82 |
| 6pm - 7pm | 25 | 41 | 34 | 24 - 51 | 41 | 34 | 23 - 53 | 51 | 46 | 35 - 63 | 55 | 52 | 37 - 70 |
| 6pm - 7pm | 30 | 40 | 34 | 24 - 50 | 38 | 33 | 23 - 47 | 49 | 46 | 35 - 60 | 54 | 51 | 39 - 67 |
| 6pm - 7pm | 35 | 36 | 31 | 22 - 44 | 34 | 29 | 21 - 42 | 45 | 42 | 32 - 55 | 49 | 46 | 35 - 60 |
| 6pm - 7pm | 40 | 33 | 29 | 21 - 40 | 30 | 26 | 19 - 37 | 41 | 39 | 30 - 50 | 45 | 42 | 32 - 55 |
| 6pm - 7pm | 45 | 31 | 27 | 20 - 38 | 27 | 24 | 18 - 34 | 38 | 36 | 28 - 46 | 41 | 39 | 30 - 51 |
| 6pm - 7pm | 50 | 29 | 26 | 19 - 35 | 25 | 22 | 16 - 31 | 36 | 35 | 27 - 44 | 38 | 36 | 27 - 46 |
| 6pm - 7pm | 55 | 27 | 25 | 18 - 33 | 24 | 21 | 16 - 29 | 34 | 33 | 26 - 42 | 35 | 33 | 25 - 43 |
| 6pm - 7pm | 60 | 26 | 24 | 18 - 32 | 24 | 21 | 15 - 28 | 32 | 30 | 24 - 39 | 34 | 31 | 24 - 41 |

**Table S3:** Typical values for frequency domain HRV features. Presented are mean, median, and the range from the 25^th^ to the 75*^th^* percentile, across gender and age ranges. Similar to Table S2, the HRV values vary significantly with time of day.

HRV Features - Frequency Domain
| Time of day | Age<br>(yr) | HF (female) |  |  | HF (male) |  |  | LF (female) |  |  | LF (male) |  |  |
| --- | --- | --- | --- | --- | --- | --- | --- | --- | --- | --- | --- | --- | --- |
|  |  | (mean) | (med) | (25p-75p) | (mean) | (med) | (25p-75p) | (mean) | (med) | (25p-75p) | (mean) | (med) | (25p-75p) |
| 6am - 7am | 20 | 1311 | 759 | 359 - 1566 | 1352 | 868 | 448 - 1665 | 1875 | 1372 | 737 - 2395 | 2265 | 1749 | 991 - 2925 |
| 6am - 7am | 25 | 971 | 543 | 250 - 1147 | 970 | 582 | 269 - 1169 | 1568 | 1120 | 570 - 2014 | 1954 | 1486 | 779 - 2551 |
| 6am - 7am | 30 | 822 | 474 | 232 - 962 | 779 | 488 | 251 - 922 | 1418 | 1018 | 547 - 1798 | 1816 | 1406 | 793 - 2341 |
| 6am - 7am | 35 | 657 | 375 | 189 - 749 | 599 | 375 | 195 - 710 | 1209 | 859 | 471 - 1530 | 1600 | 1227 | 694 - 2055 |
| 6am - 7am | 40 | 497 | 290 | 150 - 567 | 464 | 286 | 151 - 543 | 1015 | 721 | 397 - 1282 | 1389 | 1048 | 588 - 1788 |
| 6am - 7am | 45 | 379 | 225 | 118 - 432 | 366 | 221 | 118 - 411 | 842 | 593 | 331 - 1048 | 1176 | 876 | 490 - 1502 |
| 6am - 7am | 50 | 313 | 190 | 101 - 356 | 283 | 171 | 92 - 317 | 728 | 508 | 288 - 893 | 990 | 709 | 392 - 1242 |
| 6am - 7am | 55 | 280 | 170 | 91 - 314 | 245 | 138 | 75 - 255 | 659 | 450 | 254 - 794 | 853 | 586 | 321 - 1033 |
| 6am - 7am | 60 | 251 | 146 | 79 - 270 | 243 | 116 | 64 - 214 | 576 | 374 | 210 - 670 | 759 | 469 | 259 - 858 |
| 6pm - 7pm | 20 | 698 | 359 | 167 - 781 | 710 | 397 | 188 - 803 | 1071 | 733 | 394 - 1343 | 1339 | 999 | 538 - 1681 |
| 6pm - 7pm | 25 | 478 | 235 | 108 - 515 | 436 | 214 | 90 - 482 | 820 | 524 | 273 - 1015 | 1035 | 690 | 331 - 1312 |
| 6pm - 7pm | 30 | 419 | 214 | 106 - 445 | 381 | 192 | 97 - 400 | 761 | 504 | 279 - 924 | 971 | 698 | 390 - 1232 |
| 6pm - 7pm | 35 | 327 | 173 | 87 - 348 | 285 | 145 | 73 - 292 | 628 | 418 | 231 - 749 | 809 | 558 | 310 - 1012 |
| 6pm - 7pm | 40 | 254 | 142 | 75 - 270 | 216 | 112 | 58 - 225 | 524 | 351 | 199 - 612 | 675 | 457 | 246 - 831 |
| 6pm - 7pm | 45 | 213 | 119 | 63 - 223 | 172 | 93 | 48 - 175 | 440 | 293 | 167 - 508 | 557 | 375 | 202 - 673 |
| 6pm - 7pm | 50 | 184 | 104 | 55 - 192 | 136 | 74 | 39 - 139 | 383 | 254 | 145 - 446 | 455 | 302 | 164 - 540 |
| 6pm - 7pm | 55 | 160 | 90 | 49 - 165 | 129 | 63 | 34 - 116 | 334 | 221 | 125 - 380 | 386 | 244 | 131 - 437 |
| 6pm - 7pm | 60 | 143 | 79 | 44 - 145 | 135 | 57 | 32 - 106 | 292 | 186 | 107 - 325 | 358 | 204 | 110 - 373 |

**Table S4:** Typical HRV values for graphical domain Poincare *S*_1_ and *S*_2_ features. Similar to Tables S2 and S3, presented are mean, median, and the range from the 25^th^ to the 75*^th^* percentile, across gender and age ranges.

HRV Features - Graphical Domain
| Time of day | Age<br>(yr) | Poincare $S_1$ (female) | | | Poincare $S_1$ (male) | | | Poincare $S_2$ (female) | | | Poincare $S_2$ (male) | | |
| --- | --- | --- | --- | --- | --- | --- | --- | --- | --- | --- | --- | --- | --- |
|  |  | (mean) | (med) | (25p-75p) | (mean) | (med) | (25p-75p) | (mean) | (med) | (25p-75p) | (mean) | (med) | (25p-75p) |
| 6am - 7am | 20 | 50 | 43 | 27 - 66 | 56 | 51 | 34 - 72 | 120 | 115 | 84 - 148 | 141 | 134 | 103 - 171 |
| 6am - 7am | 25 | 43 | 37 | 23 - 57 | 45 | 40 | 25 - 59 | 113 | 107 | 77 - 141 | 128 | 121 | 86 - 160 |
| 6am - 7am | 30 | 40 | 35 | 22 - 52 | 40 | 36 | 23 - 53 | 112 | 106 | 77 - 137 | 125 | 116 | 83 - 154 |
| 6am - 7am | 35 | 37 | 31 | 21 - 46 | 35 | 31 | 20 - 45 | 109 | 101 | 73 - 134 | 120 | 109 | 77 - 149 |
| 6am - 7am | 40 | 33 | 28 | 19 - 41 | 31 | 27 | 18 - 40 | 104 | 96 | 70 - 127 | 115 | 104 | 72 - 143 |
| 6am - 7am | 45 | 29 | 25 | 17 - 36 | 28 | 24 | 16 - 35 | 99 | 91 | 66 - 122 | 112 | 99 | 69 - 140 |
| 6am - 7am | 50 | 27 | 23 | 16 - 33 | 25 | 21 | 15 - 30 | 98 | 89 | 64 - 122 | 105 | 92 | 64 - 131 |
| 6am - 7am | 55 | 25 | 22 | 15 - 30 | 24 | 20 | 14 - 28 | 98 | 87 | 63 - 121 | 104 | 90 | 63 - 129 |
| 6am - 7am | 60 | 24 | 21 | 15 - 29 | 25 | 19 | 14 - 27 | 95 | 84 | 60 - 118 | 103 | 89 | 62 - 131 |
| 6pm - 7pm | 20 | 37 | 31 | 20 - 46 | 39 | 32 | 20 - 49 | 90 | 83 | 61 - 109 | 102 | 93 | 70 - 127 |
| 6pm - 7pm | 25 | 31 | 25 | 17 - 37 | 30 | 23 | 15 - 37 | 82 | 74 | 54 - 100 | 90 | 81 | 59 - 113 |
| 6pm - 7pm | 30 | 30 | 24 | 17 - 36 | 29 | 24 | 16 - 34 | 81 | 74 | 54 - 100 | 91 | 84 | 62 - 110 |
| 6pm - 7pm | 35 | 27 | 23 | 16 - 32 | 26 | 21 | 14 - 30 | 77 | 70 | 52 - 93 | 83 | 76 | 55 - 103 |
| 6pm - 7pm | 40 | 24 | 21 | 15 - 28 | 22 | 18 | 13 - 26 | 71 | 64 | 48 - 87 | 76 | 69 | 50 - 94 |
| 6pm - 7pm | 45 | 23 | 20 | 14 - 27 | 20 | 17 | 12 - 23 | 68 | 62 | 46 - 83 | 72 | 65 | 47 - 88 |
| 6pm - 7pm | 50 | 21 | 18 | 13 - 25 | 18 | 15 | 11 - 21 | 64 | 58 | 44 - 77 | 67 | 60 | 45 - 83 |
| 6pm - 7pm | 55 | 20 | 17 | 13 - 23 | 18 | 14 | 11 - 20 | 63 | 57 | 42 - 76 | 64 | 57 | 43 - 79 |
| 6pm - 7pm | 60 | 19 | 16 | 12 - 22 | 18 | 14 | 10 - 19 | 61 | 54 | 41 - 74 | 61 | 55 | 40 - 75 |

**Table S5:** Sympathovagal balance: The LF/HF ratio is the estimate of the relative strengths of the sympathetic and parasympathetic branches. Presented are mean, median, and the range from the 25^th^ to the 75*^th^* percentile, across gender and age ranges.

| Time of day | Age<br>(yr) | LF / HF (female) |  |  | LF / HF (male) |  |  |
| --- | --- | --- | --- | --- | --- | --- | --- |
|  |  | (mean) | (med) | (25p-75p) | (mean) | (med) | (25p-75p) |
| 6am - 7am | 20 | 2.175 | 1.765 | 1.124 - 2.748 | 2.505 | 2.007 | 1.239 - 3.170 |
| 6am - 7am | 25 | 2.486 | 2.004 | 1.258 - 3.149 | 3.207 | 2.500 | 1.504 - 4.097 |
| 6am - 7am | 30 | 2.608 | 2.091 | 1.313 - 3.316 | 3.547 | 2.827 | 1.744 - 4.560 |
| 6am - 7am | 35 | 2.798 | 2.230 | 1.391 - 3.558 | 3.991 | 3.198 | 1.951 - 5.116 |
| 6am - 7am | 40 | 3.045 | 2.431 | 1.513 - 3.870 | 4.432 | 3.583 | 2.164 - 5.744 |
| 6am - 7am | 45 | 3.257 | 2.593 | 1.602 - 4.135 | 4.819 | 3.877 | 2.359 - 6.230 |
| 6am - 7am | 50 | 3.378 | 2.674 | 1.650 - 4.281 | 5.125 | 4.103 | 2.508 - 6.608 |
| 6am - 7am | 55 | 3.351 | 2.641 | 1.625 - 4.257 | 5.202 | 4.148 | 2.509 - 6.708 |
| 6am - 7am | 60 | 3.282 | 2.581 | 1.572 - 4.164 | 5.071 | 4.021 | 2.404 - 6.534 |
| 6pm - 7pm | 20 | 2.441 | 2.021 | 1.260 - 3.117 | 3.000 | 2.480 | 1.579 - 3.801 |
| 6pm - 7pm | 25 | 2.653 | 2.208 | 1.401 - 3.371 | 3.840 | 3.141 | 1.953 - 4.952 |
| 6pm - 7pm | 30 | 2.794 | 2.334 | 1.487 - 3.544 | 4.228 | 3.545 | 2.218 - 5.511 |
| 6pm - 7pm | 35 | 2.887 | 2.385 | 1.544 - 3.705 | 4.547 | 3.845 | 2.467 - 5.817 |
| 6pm - 7pm | 40 | 2.971 | 2.471 | 1.595 - 3.770 | 4.794 | 4.003 | 2.550 - 6.118 |
| 6pm - 7pm | 45 | 2.970 | 2.471 | 1.592 - 3.716 | 4.785 | 4.000 | 2.504 - 6.078 |
| 6pm - 7pm | 50 | 2.970 | 2.500 | 1.607 - 3.729 | 4.871 | 4.028 | 2.524 - 6.203 |
| 6pm - 7pm | 55 | 2.913 | 2.409 | 1.571 - 3.677 | 4.623 | 3.803 | 2.360 - 5.917 |
| 6pm - 7pm | 60 | 2.815 | 2.313 | 1.512 - 3.521 | 4.287 | 3.482 | 2.165 - 5.504 |

**Figure S1:**
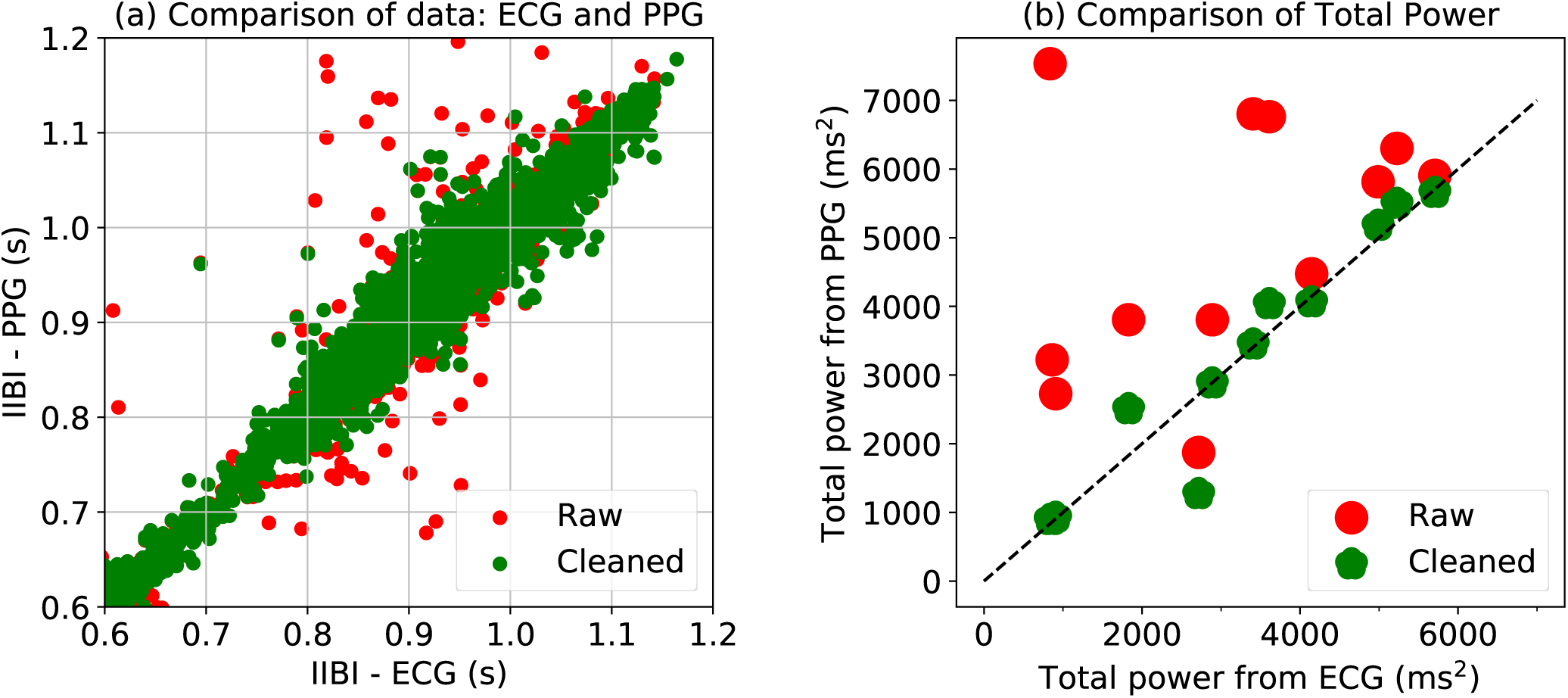
Comparison of PPG and ECG: Data is simultaneously obtained from subjects using PPG and ECG. The data is then interpolated so as to obtain data points at exactly the same time. (a) show the correlation between the ECG derived interpolated RR intervals and the PPG derived interbeat intervals (IBI). (b) shows the correlation between the total power obtained from the interpolated RR tachogram and the total power obtained from the interpolated IBI.

**Figure S2:**
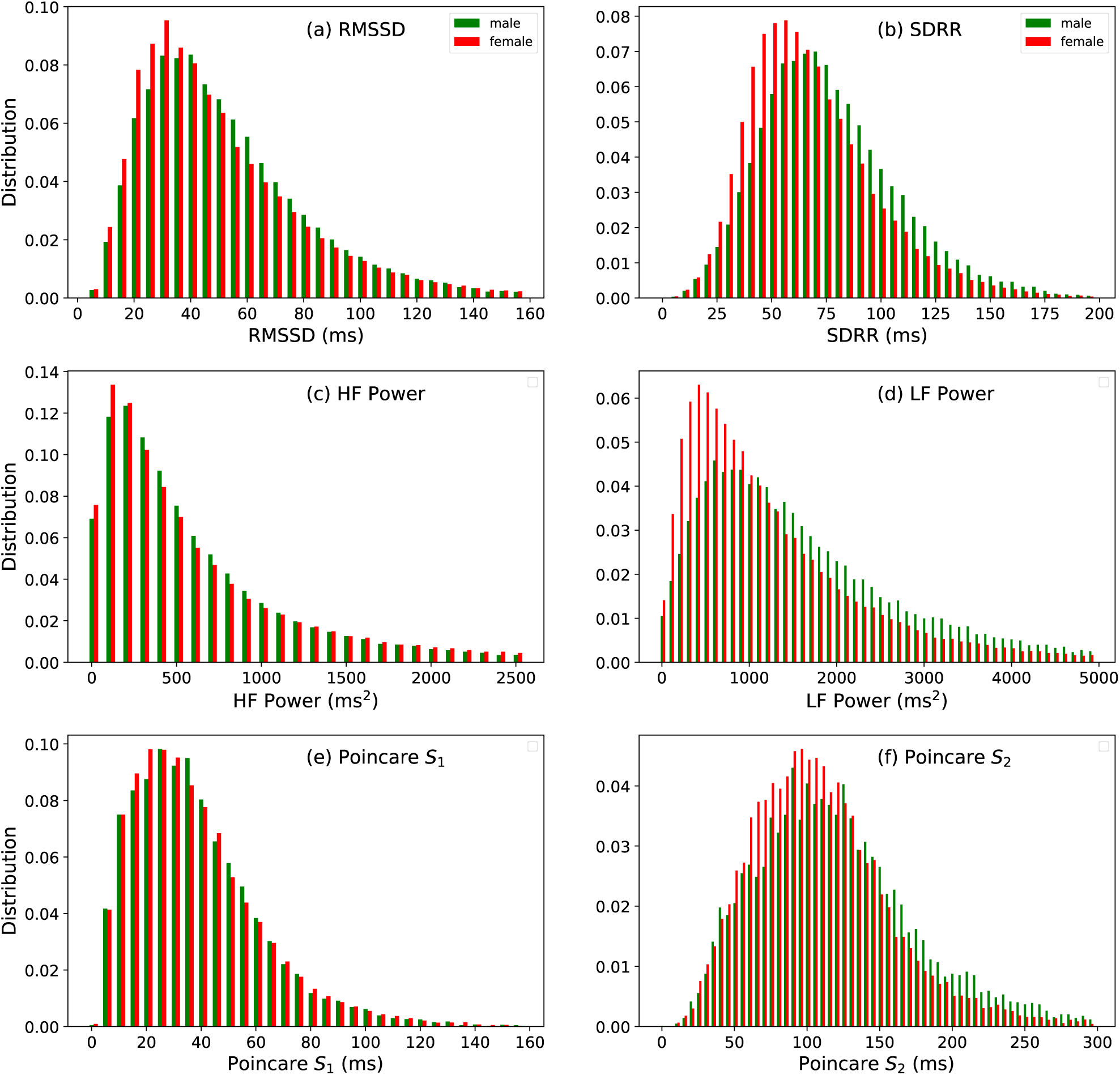
The distribution of HRV metrics for individuals in the age range 30 - 31 yr. The indicators of parasympathetic function namely, the RMSSD, HF power, and Poincare *S*_1_ distributions are similar, and show little variation with gender. On the other hand, men have higher SDRR, LF power, and Poincare *S*_2_ on average, compared to women, which indicates higher sympathetic activity.

**Figure S3:**
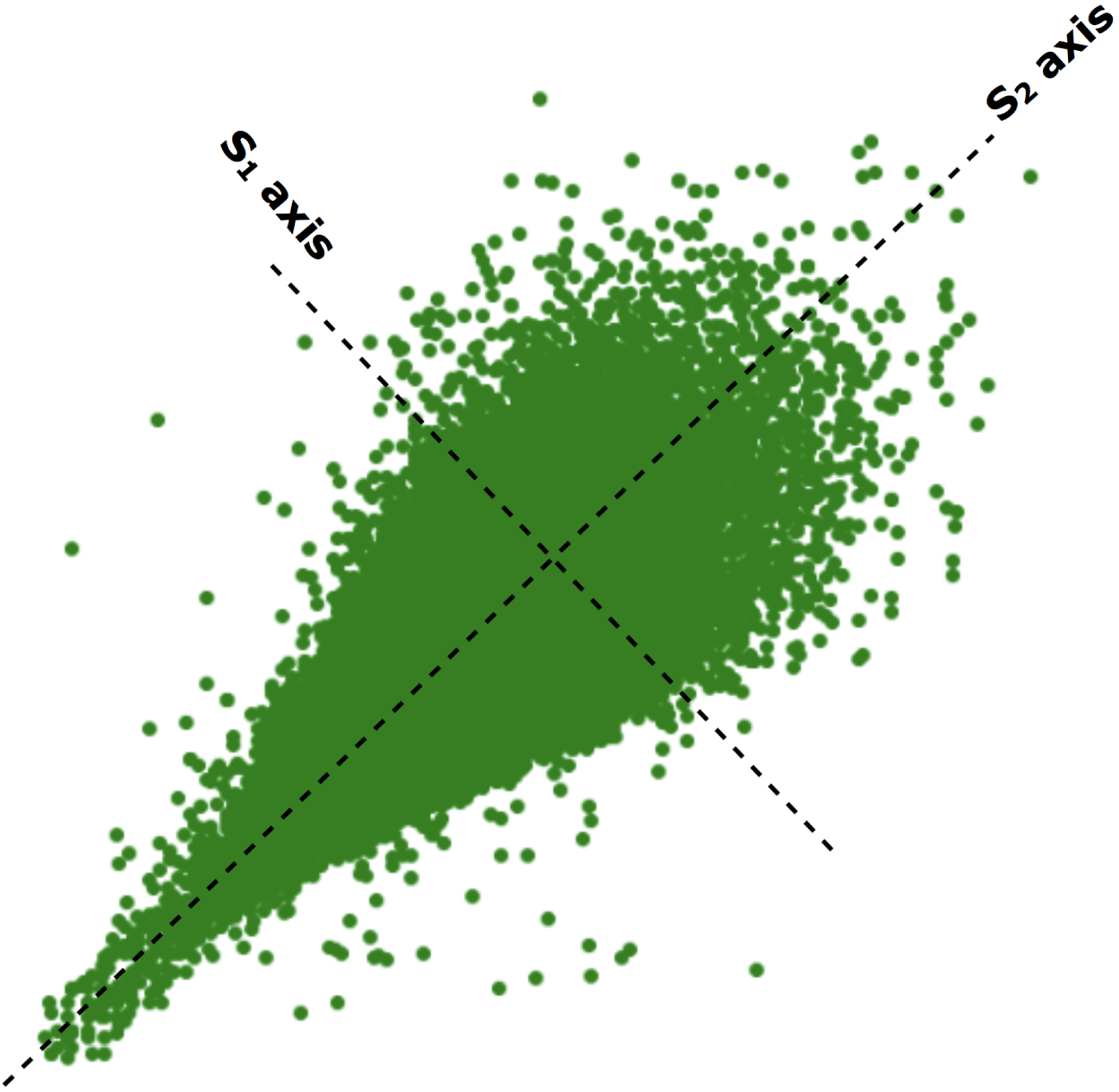
Poincare scatter plot: IBI at time index *i* plotted against the IBI at time index (*i* + 1). The Poincare plot resembles a tapered ellipse since the variability is larger for large values of IBI.

